# Conditional CAR T cells with specificity to oncofetal glycosaminoglycans in solid tumors

**DOI:** 10.1101/2024.05.29.596014

**Authors:** Nastaran Khazamipour, Htoo Zarni Oo, Nader Al-Nakouzi, Mona Marzban, Nasrin Khazamipour, Morgan E. Roberts, Negin Farivar, Igor Moskalev, Joey Lo, Fariba Ghaidi, Irina Nelepcu, Alireza Moeen, Sarah Truong, Robert Dagil, Swati Choudhary, Tobias Gustavsson, Beibei Zhai, Sabine Heitzender, Ali Salanti, Poul H Sorensen, Mads Daugaard

## Abstract

Glycosaminoglycans are often deprioritized as targets for synthetic immunotherapy due to the complexity of glyco-epitopes and limited options for obtaining specific subtype-binding. Solid tumors express proteoglycans that are modified with oncofetal chondroitin sulfate (CS), a modification normally restricted to the placenta. Here, we report the design and functionality of conditional chimeric antigen receptor (CAR) T cells with selectivity to oncofetal CS. Following expression in T cells, the CAR could be ‘armed’ with recombinant VAR2CSA lectins (rVAR2) to target tumor cells expressing oncofetal CS. While un-armed CAR T cells remained inactive in the presence of target cells, VAR2-armed CAR T cells displayed robust activation and the ability to eliminate diverse tumor cell types *in vitro*. Cytotoxicity of the CAR T cells was proportional to the concentration of rVAR2 available to the CAR, offering a potential molecular handle to finetune CAR T cell activity. *In vivo*, armed CAR T cells rapidly targeted bladder tumors and increased survival of tumor-bearing mice. Thus, our work indicates that cancer-restricted glycosaminoglycans can be exploited as potential targets for CAR T cell therapy.

## INTRODUCTION

Chondroitin sulfate (CS) glycosaminoglycans are abundantly present in the placenta where they support growth and motility of trophoblast cells (1–3). Motility is an essential feature of villous trophoblasts that allow them to invade into the uterine tissue during placental implementation (4, 5). *Plasmodium falciparum* malaria parasites express VAR2CSA lectins on the exterior of infected red blood cells, mediating unique binding-specificity to placental-type CS (6–8). In normal physiology, placental-type CS is exclusively present in the placental syncytium and VAR2CSA^+^ malaria infected erythrocytes therefore only accumulate in the placenta.

Cell growth and motility are features shared between trophoblasts and tumor cells (9–12). Perhaps to mimic the features of the placental compartment that support tumor growth, tumor cells — including those in many pediatric and adult solid tumors — re-express placental-type chondroitin sulfate (CS), as a secondary modification to a limited repertoire of proteoglycans (8, 13–17). Accordingly, recombinant VAR2CSA (rVAR2) can be used to guide therapeutic modalities towards solid tumors when formulated as rVAR2-drug conjugates (8, 15, 18, 19) or bi-specific immune cell engagers (19–21).

Oncofetal tumor antigens have emerged as potential attractive targets for chimeric antigen receptor (CAR) T cell therapy (22, 23). CARs are genetically engineered synthetic antigen receptors that are able to activate upon antigen recognition, independent of MHC presentation (24, 25). CAR T cell therapy has achieved unprecedented success in treating patients with hematopoietic malignancies, such as acute B-cell lymphoblastic leukaemia and B-cell lymphomas (23, 26–28). However, only a few studies have shown promise for CAR T cell therapy in solid tumors, such as GD2-CAR T cells for relapsed and refractory Neuroblastoma tumors (29), as well as B7H3-, GD2- and IL13RA2-CAR T cells for human Glioma (30–32). CAR T cell therapy remains challenging in solid tumors due to numerous factors including architectural heterogeneity, acquired antigen down-regulation or antigen-loss, or a lack of specific surface tumor antigens (23, 33). Hence, identification of specific antigens in solid tumors is a necessary step for extending the clinical utility of CAR T cell therapy beyond hematopoietic cancers. Until now, CAR T cell therapy has primarily focused on targeting protein antigens expressed on the surface of cancer cells. However, due to the complex challenges of CAR T cell therapy in solid tumors, there is increasing interest in exploring other types of target molecules including carbohydrates, glycolipids, and glycoproteins. For example, targeting the glycosylation component of a protein rather than the protein itself offers potential advantages. First, tumor-specific protein glycoforms can offer increased tumor selectivity and possibly limit off-target effects (34–36). Second, a specific glycosylation moiety or pattern can be present on several different proteoglycans simultaneously across cell populations, including tumor stem cells, which may overcome challenges related to heterogeneity and dormancy (19). Finally, proteins that are not normally glycosylated may be subject to disease-specific glycosylation, thereby increasing the available tumor target reservoir (8, 36–38). In this study, we utilized recombinant VAR2CSA proteins to produce CAR T cells with specificity to oncofetal CS glycosaminoglycans, broadly expressed across various solid tumor cell types.

## RESULTS

### Design and validation of a conditional CAR with selectivity to oncofetal CS glycosaminoglycans

Oncofetal CS glycosaminoglycans have been described in multiple solid tumor types including sarcoma, lymphoma, glioma, melanoma, pancreatic cancer, lung cancer, colorectal cancer, breast cancer, prostate cancer, and bladder cancer (2, 8, 13–16, 18–21, 39–41). The oncofetal CS modification can be specifically targeted by rVAR2 proteins that are currently under investigation as vehicles for therapeutic delivery (8, 15, 19–21) and as reagents in liquid biopsy diagnostic applications (14, 16, 40). Indeed, the rVAR2 proteins are specific for oncofetal CS, and binding to tumor cells can be competed with soluble CS (**Supplementary Fig. 1A**) (8, 15). In primary tumor specimens, oncofetal CS is found both in the tumor stroma and on cell membranes, and expression generally increases with tumor stage (8, 15, 39). For instance, in bladder cancer patients, oncofetal CS expression is significantly associated with advanced T stage (P=0.0231) and N stage (P=0.0114). (**Supplementary Fig. 1B-D**). Compared to bladder cancer, neuroblastoma tumors express lower amount of oncofetal CS, yet the advanced-stages tumors (II-IV) demonstrate higher expression compared to stage I tumor (**Supplementary Fig. 1E-F**). Interestingly, high levels of oncofetal CS were associated with poor survival of neuroblastoma patients (**Supplementary Fig. 1G**), which is a trend also observed in other cancer indications (15, 18).

With oncofetal CS being a broadly expressed target across solid tumor indications, we decided to explore it as a target for CAR T cell therapy. We first tested the levels of oncofetal CS on naïve and activated human T cells using rVAR2 as the detection reagent to assess the risk of potential CAR-induced T cell self-elimination. Activated and naïve T cells expressed minimal-to-undetectable levels of oncofetal CS and the expression was ∼20x lower than that detected in human bladder cancer cells (**Fig. 1A**).

**Figure 1.**
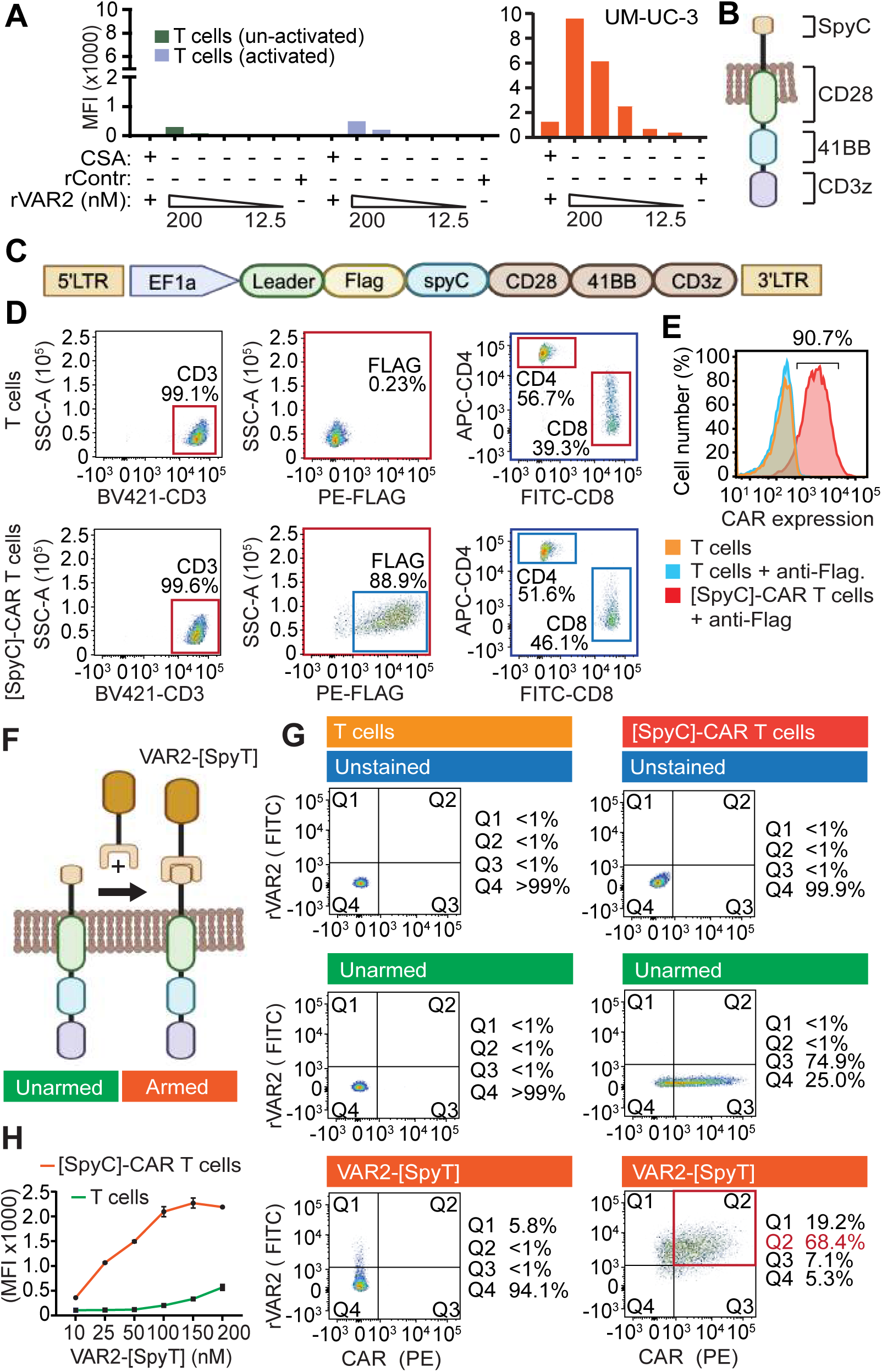
Design and validation of a conditional CAR with selectivity to oncofetal CS glycosaminoglycans. (**A**) Naïve and activated human T cells and UM-UC3 tumor cells were incubated, with control protein or different concentrations of V5-tagged VAR2 (12 - 200 nM) protein +/- purified CSA as indicated. Binding of VAR2 was assessed by flow cytometry using anti-V5-FITC. (**B**) Illustration of the [SpyC]-CAR containing intracellular CD3zeta, 41BB, and CD28 domains fused with an extracellular split-intein SpyCatcher (SpyC) domain. (**C**) Schematic diagram of the [SpyC]-CAR DNA construct in a lentiviral plasmid. (**D**) Isolated human T cells with or without [SpyC]-CAR transduction, were analyzed for CD4 and CD8 expression by flow cytometry. (**E**) [SpyC]-CAR T cells were incubated 14 days post-transduction, and expression of the [SpyC]-CAR (Flag+) was assessed by flow cytometry. The result is representative of 3 individual donors. (**F**) Schematic of [SpyC]-CAR T Cell arming with the recombinant VAR2-[SpyT] protein. (**G**) [SpyC]-CAR T cells were mixed with 200nM [SpyT]-VAR2 in triplicate and expression of Flag ([SpyC]-CAR) and V5 ([SpyT]-VAR2) were assessed by flow cytometry. (**H**) T cells with or without [SpyC]-CAR transduction were incubated with indicated concentrations of VAR2-[SpyT] protein and analyzed for VAR2-[SpyT][SpyC]-CAR assembly by flow cytometry. Error bars shown are mean ± SEM of triplicate wells. All the results are representative of 3 independent experiments.

We next designed a conditional CAR construct that utilized the specificity of rVAR2 for oncofetal CS targeting. The design allowed the CAR to be armed with rVAR2 after expression in the T cell membrane. This un-conventional design was deployed to alleviate inherent problems of expressing a functional VAR2-CAR fusion-protein in human T cells. The conditional CAR was comprised of the intracellular activation, co-stimulatory, and transmembrane domains CD3zeta, 41BB, and CD28, fused in-frame with a sequence encoding the split-protein intein domain, SpyCatcher (SpyC), derived from *Streptococcus pyogenes* fibronectin-binding protein FbaB (**Fig. 1B**) (42). For context, when the SpyCatcher domain comes into contact with the other half of the split-intein, the so-called SpyTag, they spontaneously form a covalent isopeptide bond that irreversibly links the split-intein components (42, 43). Accordingly, we anticipated this design to allow for the expression of a dormant [SpyC]-CAR in T cells that could then be subsequently armed with rVAR2 genetically fused to a SpyTag, VAR2-[SpyT]. We next transduced human T cells with the chimeric [SpyC]-CAR using lentiviral gene transfer (**Fig. 1C**). The [SpyC]-CAR sequence includes a Flag-tag that enables determination of CAR expression in T cells. Indeed, [SpyC]-CAR transduction of human T cells was ∼90% efficient and was expressed by both CD4^+^ and CD8^+^ T cell populations (**Fig. 1D**), with <10% of the population staining negative for Flag (**Fig. 1E**).

We next attempted to arm the [SpyC]-CAR T cells with the recombinant VAR2-[SpyT] warhead (**Fig. 1F**). The VAR2-[SpyT] recombinant protein contains a V5 tag that enables specific detection of armed CARs via assessment of V5 and Flag double-positive T cells. Here, 68.4% of the [SpyC]-CAR T cells could be armed with the VAR2-[SpyT] protein (**Fig. 1G**), and the spontaneous VAR2-[SpyT][SpyC]-CAR reaction on T cell membrane saturated at ∼150 nM VAR2-[SpyT] (**Fig. 1H**). In aggregate, these data demonstrate the design and expression of [SpyC]-CAR T cells that can be armed with rVAR2-[SpyT] proteins.

### VAR2-armed CAR T cells activate upon target cell engagement and produce robust cytokine responses

We next investigated whether the armed VAR2-[SpyT][SpyC]-CAR T cells became activated upon engagement with oncofetal CS-positive cancer cells. As models of adult epithelial and pediatric mesenchymal tumor types, we used UM-UC-3 muscle-invasive bladder cancer and MG-63 osteosarcoma cells, respectively, as target cell lines for most of experiments. Upon contact with UM-UC-3 and MG-63 tumor cells, VAR2-[SpyT][SpyC]-CAR T cells, but not unarmed [SpyC]-CAR T cells, upregulated the activation markers CD25 and CD69 (**Figs. 2A-B**). Similar results were obtained with LNCaP (prostate cancer) and U2OS (osteosarcoma) cell lines (**Supplementary Figs. 2A-B**). T cell activation is associated with induction of cytokines that in turn activate other immune cells, such as macrophages and dendritic cells (44). Therefore, as a secondary readout for CAR T cell activation, we examined the expression of key cytokines subsequent to VAR2-[SpyT][SpyC]-CAR T cell engagement with target tumor cells. All cell lines tested were able to trigger a robust upregulation of IFNgamma (IFNγ), IL-2, and TNFalpha (TNFα) secretion by armed VAR2-[SpyT][SpyC]-CAR T cells at orders of magnitude higher levels than that detected in unarmed [SpyC]-CAR T cells or non-transduced T cells (**Figs. 2C-D, Supplementary Figs. 2C-D**). Induction of additional cytokines such as IL-4, IL-6, and IL-13 was also detected in all the cell lines tested (**Supplementary Fig. 3**). Combined, these data show that armed VAR2-[SpyT][SpyC]-CAR T cells became activated upon target tumor cell engagement.

**Figure 2.**
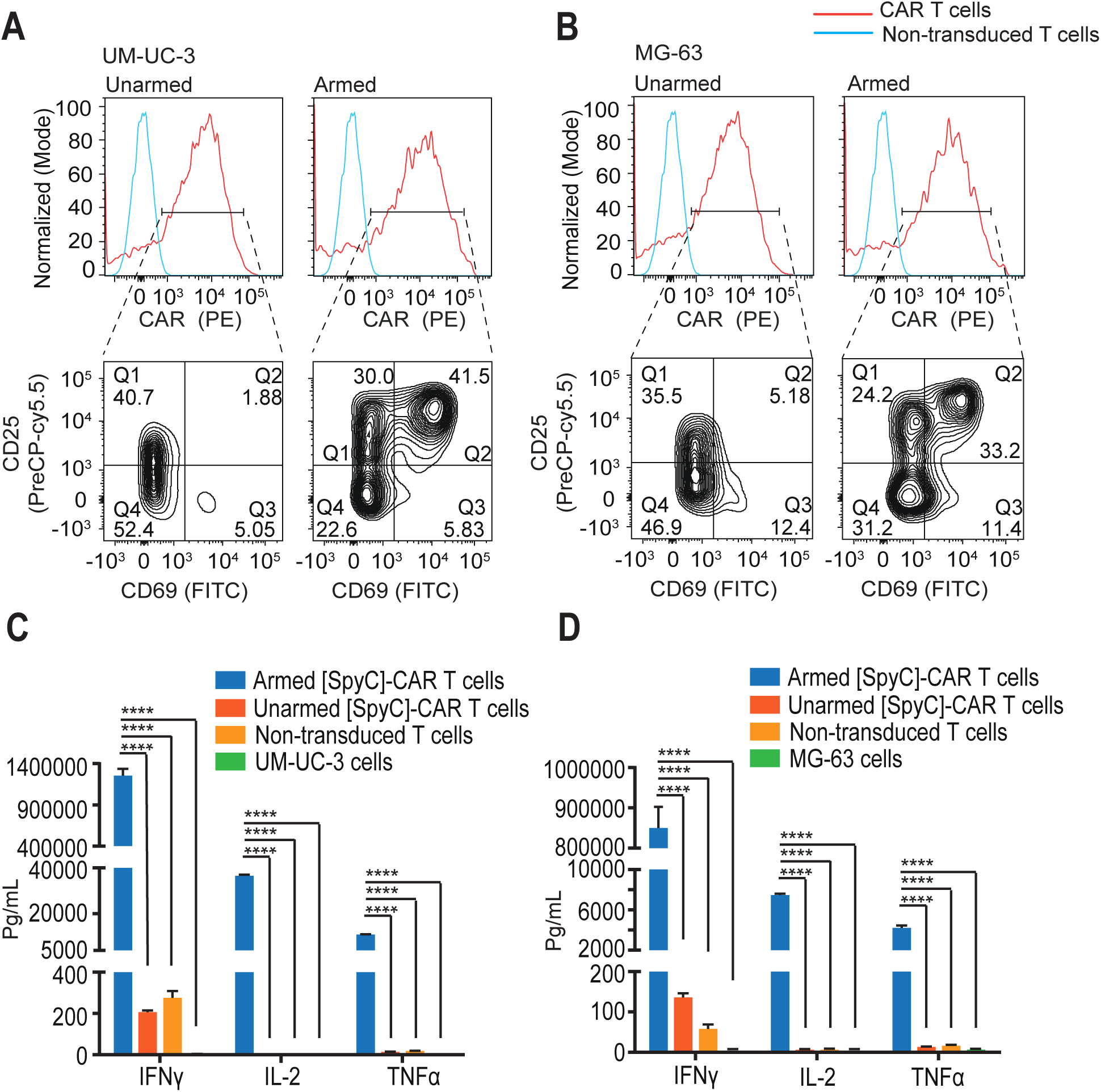
VAR2-armed CAR T cells activate upon target cell engagement and produce robust cytokine responses. (**A-B**) Armed and unarmed [SpyC]-CAR T cells were incubated, in triplicate, with (**A**) UM-UC-3 and (**B**) MG-63 cells at a 1:1 E:T ratio for 24 hours before analysis for expression of Flag, CD69, and CD25 by flow cytometry. The results are representative of 3 independent experiments. (**C-D**) Armed and unarmed [SpyC]-CAR T cells were incubated with (**C**) UM-UC-3 and (**D**) MG-63 cells at a 10:1 E:T ratio in 100 μl media, for 48 hours and concentrations of the indicated cytokines in the supernatants were quantified. Results are presented as mean ± SEM of three different wells. The statistical significance was determined using one-way ANOVA, Dunnett’s multiple comparison’s test. *, p<0.05; **, p<0.01 ***, p<0.001; ****p<0.0001.

### VAR2-[SpyT][SpyC]-CAR T cell cytotoxicity is target-cell type-dependent

We next investigated the spatiotemporal relationship between activated VAR2-[SpyT][SpyC]-CAR T cells and target cell cytotoxicity. For this, we added VAR2-[SpyT] armed and unarmed [SpyC]-CAR T cells to UM-UC-3 and MG-63 target cells at a 1:1 E:T ratio and imaged the co-cultures for 3 days. Armed VAR2-[SpyT][SpyC]-CAR T cells underwent clonal expansion (green cells) upon target cell engagement, while unarmed [SpyC]-CAR T cells remained inactive (**Fig. 3A**). Moreover, clonal expansion of the VAR2-[SpyT][SpyC]-CAR T cells efficiently eliminated the target tumor cells (red cells), while cells exposed to unarmed [SpyC]-CAR T cells outgrew the culture by day 3. Notably, VAR2-[SpyT][SpyC]-CAR T cell expansion could be detected in MG-63 cultures at day 1, while similar level of expansion was detected at day 2 in UM-UC-3 (**Fig. 3A**). To further examine potential differences in sensitivity to VAR2-[SpyT][SpyC]-CAR T cells amongst different target cell types, we subjected our diverse cell line panel to VAR2-[SpyT][SpyC]-CAR T cells in an E:T-ratio of 1:1 and recorded tumor cell viability over a week. While a decrease in MG-63 cells was observed after 18 hrs, UM-UC-3 cells needed 36 hrs of VAR2-[SpyT][SpyC]-CAR T cell exposure before a reduction in cell viability could be detected (**Fig. 3B**). LNCaP and U2OS also required 36 hrs of VAR2-[SpyT][SpyC]-CAR T cell exposure to exhibit a reduction in cell viability (**Supplementary Fig. 4A**). We next plotted the oncofetal CS expression levels of the different cell types (**Fig. 1A** and **Supplementary Fig. 1A**) against the time required for VAR2-[SpyT][SpyC]-CAR T cells to decrease viability of the same cells (**Fig. 3B** and **Supplementary Fig. 4A**). The analysis revealed that the level of oncofetal CS expression (MFI) did not directly reflect *in vitro* efficacy of VAR2-[SpyT][SpyC]-CAR T cells (**Fig. 3C**). The same was observed for oncofetal CS target cell expression and expression of key T cell activation markers (i.e., CD25^+^/CD69^+^ and IFNγ) following co-culture (**Supplementary Fig. 4C-D**). Combined, these data show that oncofetal CS-positive tumor cells of different lineages can be targeted and eliminated by VAR2-[SpyT][SpyC]-CAR T cells. The data further indicate that the different amounts of oncofetal CS expressed on the target cells all induce sufficient CAR T cell activation after target cell engagement.

**Figure 3.**
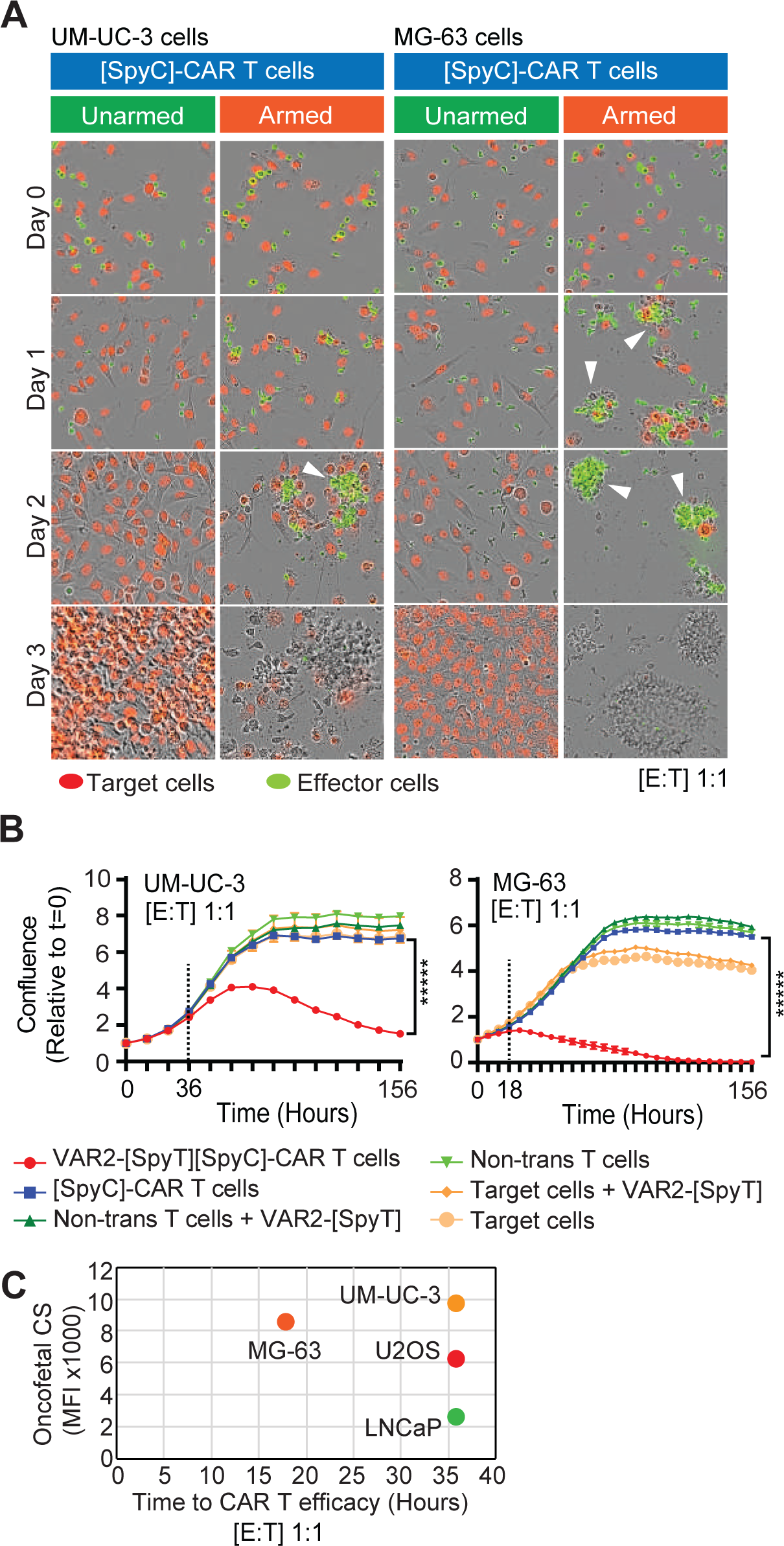
VAR2-[SpyT][SpyC]-CAR T cell cytotoxicity is target cell type-dependent and does not directly reflect oncofetal CS content. **(A)** [SpyC]-CAR T cells (green) +/- VAR2-[SpyT] protein, were co-cultured with UM-UC-3 and MG-63 target cells (red) at a 1:1 E:T ratio, in triplicate, and monitored over 3 days by IncuCyte. **(B)** Red fluorescent expressing UM-UC-3 and MG-63 target cells were co-cultured with indicated formulations of T cells and the confluency of tumor cells monitored over time by IncuCyte. Dashed lines indicate time to VAR2-[SpyT][SpyC]-CAR T cell cytotoxicity. Error bars represent mean ± SEM of triplicate wells. Data from a representative donor of three individual donors is shown. Statistics were calculated for the last timepoint, using one-way ANOVA and Dunnett’s multiple comparisons test (**C**) Oncofetal CS expression levels of different cell types were plotted against the time required for VAR2-[SpyT][SpyC]-CAR T cells to induce cytotoxicity.

### [SpyC]-CAR T cells can be conditionally armed with VAR2-[SpyT] to eliminate tumor cells

The [SpyC]-CAR was designed to allow subsequent and conditional arming of [SpyC]-CAR T cells using rVAR2-[SpyT] proteins in a dose dependent manner **(Figs. 4A)**. To substantiate our finding, we evaluated [SpyC]-CAR T cells cytotoxicity to UM-UC-3 and MG-63 target cells after exposure to increasing concentrations of VAR2-[SpyT] (0-200 nM) (**Fig. 4A**). We observed a concentration-dependent decrease in target cell viability over the week after CAR T cell exposure that reflected the number of VAR2-[SpyT] proteins available for the [SpyC]-CAR T cells during arming (**Fig. 4B**). A similar trend was observed in LNCaP and U2OS cells (**Supplementary Fig. 4B**). Notably, UM-UC-3 cells seemed slightly less sensitive to VAR2-[SpyT][SpyC]-CAR T cells in the 25-200 nM VAR2-[SpyT] concentration range as compared to MG-63 (**Fig. 4B**). To further characterize the difference in sensitivity between the target cell lines, we examined VAR2-[SpyT][SpyC]-CAR T cell cytotoxicity in various effector-to-target cell (E:T) ratios. As expected, higher CAR T cell effectors relative to target cells resulted in greater cytotoxicity; however, UM-UC-3 cells generally required more effector cells than MG-63 (**Fig. 4C**), further reflecting the less sensitivity of UM-UC-3 to VAR2-[SpyT][SpyC]-CAR T cells (**Fig. 4B-C**). Combined, these data show that [SpyC]-CAR T cells can be conditionally armed with the VAR2-[SpyT] warhead to confer an E:T ratio-dependent cytotoxicity towards target cells.

**Figure 4.**
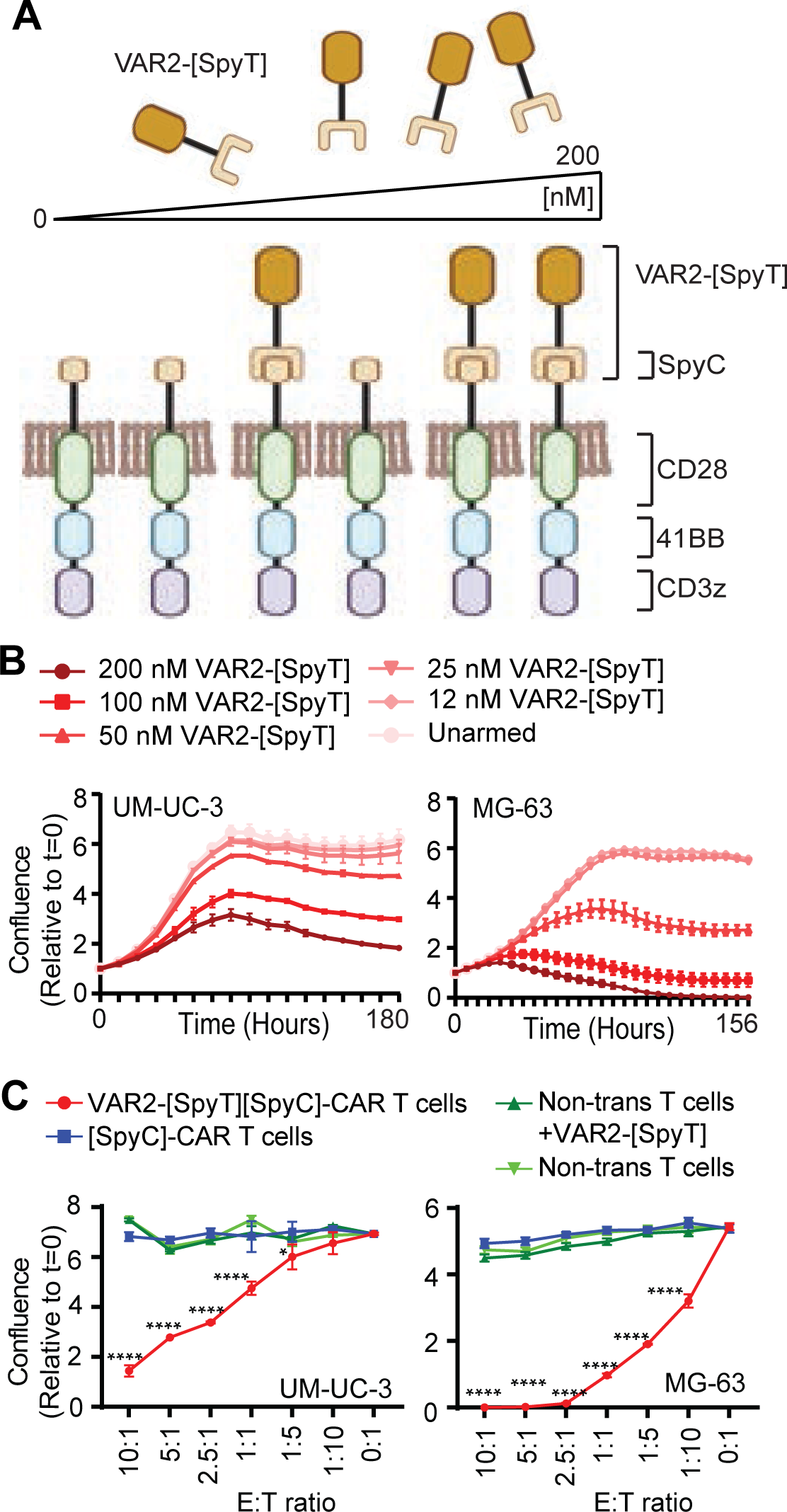
[SpyC]-CAR T cells can be conditionally armed with VAR2-[SpyT] to eliminate tumor cells. **(A**) Illustration of the conditional arming of [SpyC]-CAR T cells with increasing amounts of VAR2-[SpyT] protein. (**B**) UM-UC-3 and MG-63 target cells (red) were co-cultured with [SpyC]-CAR T cells at a 1:1 E:T ratio in triplicate, with the indicated concentrations of VAR2-[SpyT] protein, confluence was assessed as readout of tumor cell viability. Error bars represent mean ± SEM of triplicates. (**C**) UM-UC-3 and MG-63 target cells (red) were co-cultured with [SpyC]-CAR T cells +/- VAR2-[SpyT] in indicated E:T ratios and analyzed as in (B). Results are presented as mean ± SEM triplicates. Statistics were calculated, using two-way ANOVA and Dunnett’s multiple comparisons test. Data from a representative of three from 3 individual donors is shown.

### VAR2-[SpyT][SpyC]-CAR T cells inhibit tumor growth *in vivo* and prolong animal survival

CAR T cell therapy is challenging in solid tumors due to lack of durable efficacy (23, 33). Improvement in efficacy has been pursued by a variety of approaches including co-targeting of immune evasion mechanisms, combining different co-stimulatory domains in the CAR, the use of vaccines that target the CAR epitope, and the use of repeated treatments with short-lived CAR T cells (45). However, none of these approaches have resulted in sufficient increases in efficacy to date. We tested the performance of the VAR2-[SpyT][SpyC]-CAR T cells in a solid tumor xenograft mouse model using UM-UC-3 bladder tumor cells. Since the half-life of the VAR2 protein in blood circulation is very short, it is challenging to obtain sufficient exposure to the newly proliferated [SpyC]-CAR T cells in mice. Therefore, instead of injecting additional doses of VAR2-[SpyT] protein to arm the [SpyC]-CARs during clonal expansion, we decided to arm the [SpyC]-CAR T cells *in vitro,* and then inject several doses of armed CARs into mice. Nude mice were inoculated subcutaneously with 1×10^6^ UM-UC-3 cells in their right flank (day 0). At day 7, the mice were ranked by tumor size and evenly distributed into three groups with nine mice per group. Mice were injected intravenously with PBS (Group 1), unarmed [SpyC]-CAR T cells (Group 2), or armed VAR2-[SpyT][SpyC]-CAR T cells (Group 3) on days 7, 10, 13, 16, and 19 (**Fig 5A**). After the first three injections (day 13), the VAR2-[SpyT][SpyC]-CAR T cells (Group 3) started to reduce tumor growth, which became more pronounced over time (**Fig. 5B**) and was statistically significant as compared to unarmed [SpyC]-CAR T cells (Group 2) (**Fig. 5C and Supplementary Fig. S5**). No difference was observed in tumor growth between control groups 1 and 2, indicating no unintentional targeting of tumor cells by unarmed [SpyC]-CAR T cells (Group 2) (**Fig. 5C**). In the group treated with VAR2-[SpyT][SpyC]-CAR T cells (Group 3), one mouse was tumor-free at humane endpoint of Group 1 and 2 mice, while the remaining mice had various degrees of treatment benefits as compared to the control groups. This translated into increased overall survival of mice treated with VAR2-[SpyT][SpyC]-CAR T cells (**Fig. 5D**). In summary, these data demonstrate a moderate yet significant effect of VAR2-[SpyT][SpyC]-CAR T cells in the treatment of bladder tumors *in vivo*.

**Figure 5.**
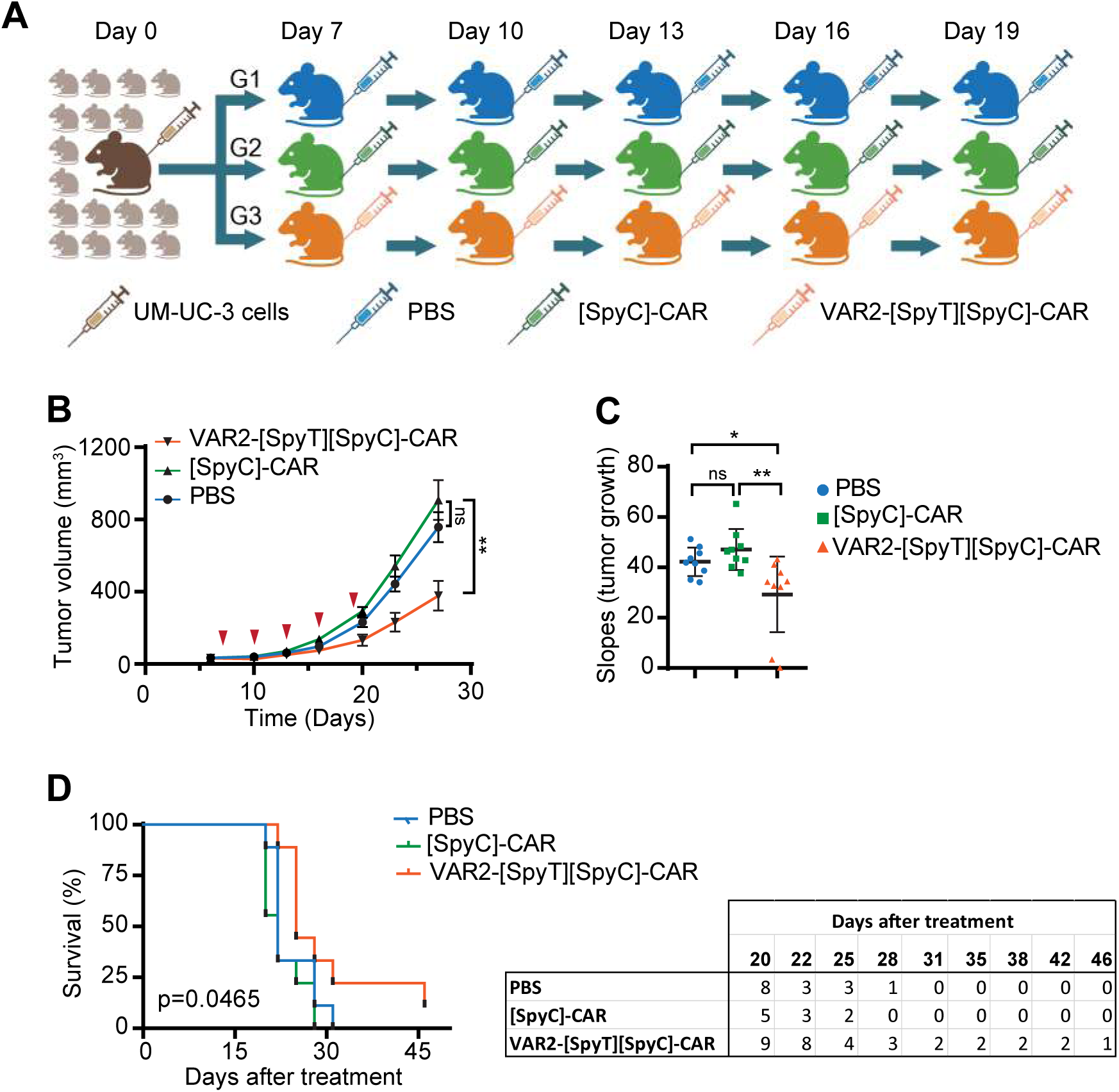
VAR2-CAR T cells suppress solid tumor growth in vivo and prolong animal survival. (**A**) Schematic outline of the experimental design. Nude mice were inoculated with 1×10^6^ UM-UC-3 tumor cells in their right flank. At day seven the mice were randomized into three groups (n=9). The groups subsequently received five doses of either PBS (G1), 2.5 million [SpyC]-CAR T cells (G2), and 2.5 million VAR2-[SpyT][SpyC]-CAR T cells (G3) by intravenous injection at the indicated times points post-tumor cell inoculation. (**B**) Tumor growth curves with the mean tumor volumes of mice in each group. Data presented as mean ± SEM (n = 9 mice). (**C**) The slope of tumor growth curve for each individual mouse in each group was calculated using linear regression, and the tumor growth rates was compared between treatment groups. The statistical analysis was performed using ANOVA followed by a Tukey’s multiple comparison’s test. (**D**) Kaplan–Meier plot (*left*) indicates morbidity as a proxy for overall survival of mice (numbers of mice) over time (*right*).

## DISCUSSION

The malarian rVAR2 protein has remarkably high specificity and affinity for oncofetal CS, which is expressed almost exclusively in the placenta and tumors (8). Therefore, targeting oncofetal CS with therapeutic formulations of the VAR2 protein has potential positive implications for cancer therapy. Oncofetal CS is present in both early and more advanced disease stages, regardless of tumor origin (15) (18). The concurrent presence of oncofetal CS across different proteins expressed by tumor cells increases the molecular density of the involved glycosaminoglycan and the potential sensitivity to oncofetal CS targeting technologies (8, 13).

Preclinical studies in cancer and early-phase clinical trials in malaria, have demonstrated the feasibility and safety of VAR2 as a therapeutic agent. A phase I clinical vaccine trial (PAMVAC) in pregnancy-associated malaria demonstrated low immunogenicity of VAR2 in humans when administrated without adjuvants, and acceptable safety profiles (46). In consideration to this subject, VAR2CSA may have co-evolved with humans to exhibit minimal immunogenicity as evidenced by the persistence of malaria endemics today. Moreover, VAR2 in formulations as drug-conjugates or bi-specific immune cell engagers (CD3-fusion proteins) all exhibit strong efficacies in both immunocompetent and immunodeficient mouse models with no off-target or immune-related side effects, as well as no organ toxicity (8, 15, 21).

Since oncofetal CS is widely expressed in solid tumors and the VAR2 protein has been credentialized as a therapeutic vehicle for drug-delivery and immune cell engagement, we proposed that VAR2-directed CAR T cells might show efficacy in solid tumors. To test this idea functionally, and to alleviate issues around the safety of current CAR therapies, we designed a conditional CAR construct that could be armed with a recombinant VAR2 protein produced in bacteria after its expression in T cell plasma membranes. We used the well-defined SpyCatcher-SpyTag intein system derived from *S. pyogenes* (42) as a molecular glue to link bacterial-produced recombinant VAR2-[SpyT] with [SpyC]-CARs translated and expressed in primary human T cells. [SpyC]-CAR T cells are indolent as they cannot engage target cells and therefore fail to activate. For activation, the CAR T cells rely on being armed with a SpyTagged binder that can be any molecule (*e.g.,* proteins, peptides, or an scFv) with specificity to a tumor-selective epitope. We used VAR2-[SpyT] protein as the warhead but in principle the [SpyC]-CAR T cells could be armed with any binder of choice. Armed VAR2-[SpyT][SpyC]-CAR T cells were able to fully activate upon engagement with oncofetal CS and to eliminate target tumor cells *in vitro* in a time and dose dependant manner that depended on availability of the VAR2-[SpyT] warhead. In a murine xenograft model of bladder tumor, VAR2-[SpyT][SpyC]-CAR T cells were able to curb tumor growth and prolong survival of the mice. However, with only one complete responder, it is clear that our strategy faces similar obstacles as other CAR T approaches in solid tumors that limits CAR T cell efficacy, and further work is required to optimize this approach.

Upon activation induced proliferation of the VAR2-[SpyT][SpyC]-CAR T, the daughter cells maintain expression of the [SpyC]-CAR but become unarmed as a result of lack of the warhead during proliferation. The cytotoxicity observed therefore comes from the parental VAR2-[SpyT][SpyC]-CAR T cells after initial target engagement. This implies that CAR T cell cytotoxicity could potentially be amplified by supplementing excess amounts of VAR2-[SpyT] protein (*i.e.,* re-arming) during the clonal expansion phase *in vivo*. This provides a safety measure allowing control over CAR T cell activation based on the availability of VAR2-[SpyT] protein. However, the strategy is limited *in vivo* by the fact that the serum half-life of rVAR2 proteins is <10 min, limiting tumor exposure to the point where the concentration of available VAR2-[SpyT] is likely insufficient for saturating newly proliferated [SpyC]-CAR T cells. *In vivo* re-arming may be possible with formulations that increases rVAR2 plasma half-life. We also observed that, the degree of oncofetal CS expression on different tumor cells alone does not appear to determine the activity and cytotoxicity of the VAR2-[SpyT][SpyC]-CAR T cells. This indicates that factors beyond pure target expression might contribute to determining tumor cell sensitivity to CAR T cells. The reasons for this observation can be many, however, a simple explanation could be intrinsic differences amongst the cell lines for sensitivity to T cell-mediated cell death. Additional research in this topic is warranted.

In summary, we have provided proof-of-concept for a conditional CAR T cell approach that can target an oncofetal glycosaminoglycan modification in solid tumors. Adding cancer-specific glycosaminoglycan modifications to the CAR T cell target repertoire provides additional opportunities for immunotherapy in solid tumors.

## MATERIAL and METHODS

### Cell lines

The cell lines used in this study were maintained in specific culture media tailored to their requirements: T cells were cultured in ImmunoCult XF T cell Expansion Medium supplemented with 100U IL2, UM-UC3 cells were cultured in MEM media supplemented with 1X non-essential amino acids, MG-63 cells were cultured in MEM, LNCaP cells were cultured in RPMI, and U2OS cells were cultured in DMEM. All media were supplemented with 10% fetal bovine serum (FBS). Cultures were maintained at 37°C incubator with 5% CO2. Regular testing for mycoplasma contamination was performed, and the cell lines were confirmed to be mycoplasma-free prior to use in experiments.

To establish stable cell lines expressing nuclear-restricted mKate2 (a far-red fluorescent protein), the NucLight Lentivirus Reagent (Essen bioscience, Cat. 4476) was utilized for transduction. Cells were seeded in 24-well plates 24 hours prior to transduction. The IncuCyte NucLight Lentivirus Reagent was added at a MOI of 3 (= TU/cell), supplemented with 10 μg/mL protamine sulfate. The plate was incubated at 37°C, 5% CO2 for 24 hours. Media was replaced the next day. After 48 hours, cells were treated with zeocin selection marker (Gibco, Cat. R25001) to select for transduced cells.

### Blood samples and T cell preparation

Leukopak from healthy donors was purchased from StemCell Technologies. T cells were separated by negative selection using EasySep™ Human T Cell Isolation Kit (StemCell, Cat. 17951). The leukopak samples were prepared by adding an equivalent volume of PBS2 and centrifuging at 500 x g for 10 minutes at room temperature (15 - 25°C) and removing the supernatant. The cells were resuspended to a concentration of 5 x 10^7^ cells/mL in PBS2. T cells were isolated according to the manufacturer. Briefly, the samples were transferred to the polystyrene round-bottom tube, 50 µL/mL EasySep™ Human T Cell Isolation Cocktail and 40 µL/mL of EasySep™ Dextran RapidSpheres™ were added to the samples and incubated for 5 min at RT. The tube was topped up by PBS2, placed in a magnet (StemCell, Cat.18001), and incubated for 3 min. By inverting the magnet, the enriched T cell suspension was transferred into a clean 14 ml tube. The cells were centrifuged and resuspended in 1 mL ImmunoCult™-XF T Cell Expansion medium and counted to adjust to the concentration to 1×10^6^ cells/mL. The cells were activated by incubating with 25 µL/mL of human CD3/CD28/CD2 T Cell activator (StemCell, Cat. 10990) for 3 days at 37°C with 5% CO_2_, and then maintained in media containing 100 U/mL IL-2 (StemCell, Cat. 78036.1)

The isolated cells were assessed for T cells purity and CD4 and CD8 sub-population proportions before and after virus transduction. T cells were stained with BV421-conjugated anti-CD3 (Clone SK7), APC-anti CD4 (Clone OKT4), FITC anti CD8 (Clone SK1), and PE-FLAG (Clone L5) antibodies for 40 minutes, washed and subjected to flow cytometry analysis. Unstained and single-color controls were acquired and used for compensation.

### Protein production

Recombinant *P. falciparum* VAR2CSA proteins were produced in *E. coli* SHuffle cells (NEB) and comprised of the minimal CS-binding region (subunit ID1-ID2a) with a C-terminal V5 tag and 6x-His tag, and an N-terminal SpyTag (8). The recombinant control (rContr) protein is made from the non-CS binding region (DBL4) of VAR2CSA protein with the addition of the C-terminal V5 tag (as in (8)).

### VAR2 binding assay

Tumor cells were cultured in their appropriate media to reach 70%–80% confluency. To minimize the risk of damage to proteoglycans, the cells were prepared for binding assay using a non-enzymatic cell dissociation solution (Cellstripper, Cat. CA45000-668). For the preparation of T cells, blood-derived T cells were isolated and divided into two parts. The first part was maintained in culture media without T cell activators, as the non-activated T cells. The second group was activated in media supplemented with CD2/CD3/CD28 T cell activator (StemCell, Cat. 10990) (25 μL/mL) and IL-2 (100 U/mL) for a duration of three days.

Prior to incubation with VAR2, the cells were washed with PBS containing 2% FBS (PBS2), centrifuged at 350 x g for 5 minutes, and resuspended to a concentration of 1×10^6^ cells/mL. 100 μl of the cell suspension was added per well in a 96 well plate, followed by centrifugation to pellet the cells. The supernatant was aspirated, and the cells were resuspended in the protein solutions. The protein solutions were prepared by serial dilutions of rVAR2 diluted in PBS2 (12.5-200 nM). A cell sample without rVAR2 was used as background for antibody signal. 200 nM rContr protein, which is a recombinant non-CS binding region (DBL4) of the full-length VAR2CSA protein (8), was used as a negative control. Binding specificity was tested by the inclusion of a high concentration (400 µg/ml) of purified CSA (Sigma Cat. 27040) which competes for VAR2 binding. For this competition, 200 nM rVAR2 was pre-incubated with CSA before adding to the cells. The plate was incubated for 30 min at 4°C on a shaker. After incubation, the cells were washed twice with PBS2 and then stained for 40 min on ice with an anti-V5 antibody conjugated to FITC (Invitrogen, Cat. R963-25). The cells were washed 3 times with 200 μl FACS buffer (PBS containing 2% FBS, 2.5 mM EDTA, and 0.05 mM NaN3) and resuspended in FACS buffer containing DAPI (0.1 µg/mL) for gating out the dead cells. Samples were acquired on a FACS Canto II flow cytometer, (BD Biosciences), and the data was analyzed using FlowJo V10.4.2. Unstained and single-color controls were acquired and used for compensation

### Generation of CAR T cells

Lentivirus particles were generated in HEK293T cells following transfection with the CAR plasmid along with pMD2.G envelope (Addgene, Cat.12259) and psPAX2 packaging plasmids (Addgene, Cat.12260), using extreme Xp transfection reagent. 5×10^6^ HEK293T cells were seeded in 10 mL of DMEM media in 10 cm poly-L-lysine coated culture plates. The next day, the media was replaced with 7 mL of fresh media, 3-4 hours before transfection. A total of 25 μg of plasmid DNA, consisting of 10 μg library plasmid, 10 μg envelope plasmid (pMD2.G) and 5 μg packaging plasmid (psPAX2), were diluted in 500 μL serum-free Opti-MEM (Cat.11058021) in a sterile 1.5 mL tube. X-tremeGENE HP DNA Transfection Reagent (Cat. 6366236001) was added in a 3:1 ratio of reagent to DNA, mixed by pipetting and incubated at room temperature for 25 min. The transfection complex was added to cells dropwise while swirling the plate, and the plate was placed in a CO2 incubator. Viral supernatant was collected 48 h and 72 h post transfection and centrifuged at 350 x g for 10 minutes to remove any cells and debris. The supernatant was filtered through an 0.45 μm Steriflip and used on the same day or stored at - 80°C for future use.

To transduce T cells, the isolated human T cells were centrifuged and resuspended in virus supernatant to a concentration of 1×10^6^ cells/mL in the presence of 10 μg/mL protamine sulfate. 72 hours after transduction, 2 µg/mL puromycin was added to the media to select for CAR expressing T cells.

Two weeks after transduction, the CAR expression levels were quantified by flow cytometry with fluorescent-conjugated Flag antibody. Dead cells were excluded by DAPI staining. The data was analyzed using the FlowJo software to calculate the percentage of the live, single cells with CAR expression.

### VAR2-[SpyT][SpyC]-CAR arming on the T cell membrane

To assess the optimal quantity of VAR2-[SpyT] required to arm the [SpyC]-CARs on the T cell membrane, a saturation study was performed. Equal numbers of [SpyC]-CAR T cells (1×10^5^) were incubated with different concentrations of VAR2-[SpyT] from 0 to 200 nM. The cells were washed three times with PBS2 and stained with FITC-conjugated anti-V5 antibody (at dilution of 1:500) and PE conjugated anti Flag antibody (at dilution of 1:100) for 45 min on ice. After staining and washing, cells were resuspended in DAPI-containing FACS buffer for flow cytometry data acquisition. The FlowJo software was utilized for data analysis, initially gating live single cells for CAR expression. Subsequently, the geometric mean fluorescent intensity (MFI) of FITC within the CAR-expressing population was determined to construct the saturation curve. Non-transduced cells were incubated with equivalent amounts of VAR2-[SpyT] and analyzed as the control group. VAR2-[SpyT] arming of the [SpyC]-CAR was also analysed on flowcytometry by assessing the co-localization of the Flag-tag in CAR construct with the V5-tag on VAR2-[SpyT], following the incubation of the cells with either 200 nM of VAR2-[SpyT] or without VAR2.

### Detection of T cell activation markers by flow cytometry

The expression of T cell activation markers, CD25 and CD69, was assessed on effector cells subsequent to encountering their targets. 1×10^5^ tumor cells were co-cultured with 1×10^5^ [SpyC]-CAR T cells, in presence (armed CAR) or absence (unarmed CAR) of VAR2-[SpyT]. All samples were prepared in triplicates. The plates were incubated at 37°C overnight, then only the suspension cells were harvested, washed, and stained with PE-conjugated anti-Flag (Clone L5), PerCP/Cy5.5-conjugated anti CD25 (Clone BC96, BioLegend, Cat. 302626), and FITC-conjugated CD69 (Clone FN50, BioLegend, Cat. 310904) antibodies for 30 min on ice. Unstained and single-color controls were acquired and used for compensation. Following three washes, the samples were resuspended in FACS buffer containing DAPI to exclude the dead cells during subsequent flow cytometry analysis. Unstained and single-color controls were utilized for compensation purpose during analysis using FlowJo software. The cells were first gated for viable single cells that were Flag positive, indicating CAR-expressing cells, and then quantified for up-regulation of CD69 and CD25.

### Cytokine analysis

Cytokine production was evaluated using MSD V-plex Proinflammatory Panel 1 Human kit (Meso Scale Discovery, Cat. K15049D) according to the manufacturer’s instruction. Effector and target cells were cocultured at an E:T ratio of 10:1 in 100 μl media in a 96 well plate and incubated for 48 hours. The CAR was armed with 200nM of VAR2-[SpyT]. Post-incubation, media from each well was collected, centrifuged at 350 x g for 5 min at 4°C to remove cells and debris. Supernatants were collected stored in -80 freezer, and then evaluated for cytokine levels according to the manufacturer’s protocol. Data analyses were performed using Discovery Workbench software.

### *In vitro* cytotoxicity assay

Red fluorescent tumor cells (mKate2^+^) were co-cultured with [SpyC]-CAR in triplicate at an E:T ratio of 1:1 in the presence or absence of VAR2-[SpyT]. Similar co-cultures involving non-transduced T cell, as well as the same number of tumor cells without effector cells were used as controls. The plates were placed in the IncuCyte S3 instrument for scanning every 6 hours for up to 7 days. At each time point, 5 images per well at 10X magnification were collected. Total red area (um2/well) was quantified as a measure of live tumor cells and values were normalized to t=0 measurement. For tracking co-localization of the effector cells with target cells, [SpyC]-CAR T cells were labelled with IncuCyte® CytoLight Rapid Green Reagent. These green-labelled T cells were then co-cultured with red fluorescent tumor cells at a 1:1 ratio in 96 well plates, in the presence or absence of VAR2-[SpyT], and scanned by IncuCyte at 20X magnification for 3 days.

### In vivo model

Female nude mice (8 weeks old; Jackson Laboratory) were subcutaneously inoculated in their right flanks with 1×10^6^ UM-UC-3 cells suspended in PBS and Matrigel® Matrix solution (1:1). At day seven post-inoculation, mice were evenly distributed into three groups by tumor size, with nine mice allocated to each group. The first group received five intravenous (IV) injection of PBS, the second group received five doses of 2.5 million [SpyC]-CAR T cells, and the third group was injected with five doses of 2.5 million VAR2-[SpyT][SpyC]-CAR T cells. The tumor sizes were measured twice per week using a caliper, and tumor volume was calculated by V= W * L * H * pi/6 formula, where w=width of tumor, L=length of tumor and H=height of the tumor. The mice were euthanized when reaching tumor size of 1000LJmm3. Statistical analysis was performed using GraphPad Prism (GraphPad Software). Two-way ANOVA using Tukey’s multiple comparison tests was applied on tumor growth data. Data were presented as mean ± SEM. To quantify the rate of tumor growth over time, we calculated the slope of the tumor growth curve for each individual mouse in each group using linear regression (**Figs. 5C, Supplementary Figs. 5)**. The statistical significance of the differences between groups was assessed using one-way ANOVA, followed by Tukey’s multiple comparison test. In order to compare the survival rates between groups, Kaplan-Meier survival curve was generated, utilizing the morbidity of mice as a surrogate for overall survival across time.

## Author contributions

Conceptualization, Nt.K., N.AN, A.S., P.S., SH and M.D.; Methodology, Nt.K., H.Z.O., M.M., N.F., M.E.R., A.M., B.Z., I.M., J.L., F.G., I.N., T.G., S.C., R.D.; Data acquisition and analysis, Nt.K., H.Z.O., S.T., Nr.K., Funding acquisition, M.D., P.S., A.S.; First manuscript draft, Nt.K., S.T., P.S., and M.D., Review & Editing, all authors; Supervision, M.D., N.AN, A.S.

## Acknowledgements

We thank Drs. Sabine Heitzeneder and Crystal Mackall (Stanford, CA) for intellectual input on CAR T cell methodology, and all members of the Daugaard Lab for helpful discussions. We thank the funding support from NIH Prostate Cancer PNW-SPORE (1016339, 223493; 5P50 CA097186-17); Canadian Institutes of Health Research (CIHR) (PJT-153092); St. Baldrick’s Foundation/American Association for Cancer Research/Stand Up to Cancer Pediatric Dream Team Translational Research Grant (to PHS and MD; SU2C-AACR-DT-27-17). Stand Up to Cancer (SU2C) is a division of the **<u>Entertainment Industry Foundation</u>**, and research grants are administered by the **<u>American Association for Cancer Research</u>**, the scientific partner of SU2C. Nastaran Khazamipour (Nt.K) was supported by the CIHR-Vanier scholarship (#01353-000), UBC Four Year Fellowships (FYF) (#6456), Canadian Urological Association Scholarship Foundation (CUASF-BCC) Research Grant, and the CIHR travel Award-Michael Smith Foreign Study Supplement (#6580) for an internship with Stanford University (CA). AS is supported by NNF Tandem grant (NNF21OC0068192) and NNF Distinguished Innovator grant (NNF22OC0076055);

## Conflicts of Interest

M.D. as the corresponding author certifies that all conflicts of interest, including specific financial interests and relationships and affiliations relevant to the subject matter or materials discussed in the manuscript (e.g., employment/affiliation, grants or funding, consultancies, honoraria, stock ownership or options, expert testimony, royalties, or patents filed, received, or pending), are the following: M.D., A.S., and P.H.S. are co-founders of, and shareholders in, VAR2 Pharmaceuticals. N.A.N., and T.G. are consultants for VAR2 Pharmaceuticals. VAR2 Pharmaceuticals is a biotechnology company that specializes in therapeutic development of the VAR2CSA technology (www.var2pharma.com). The remaining authors declare no conflicts of interest.

## Supplementary

**Figure S1.**
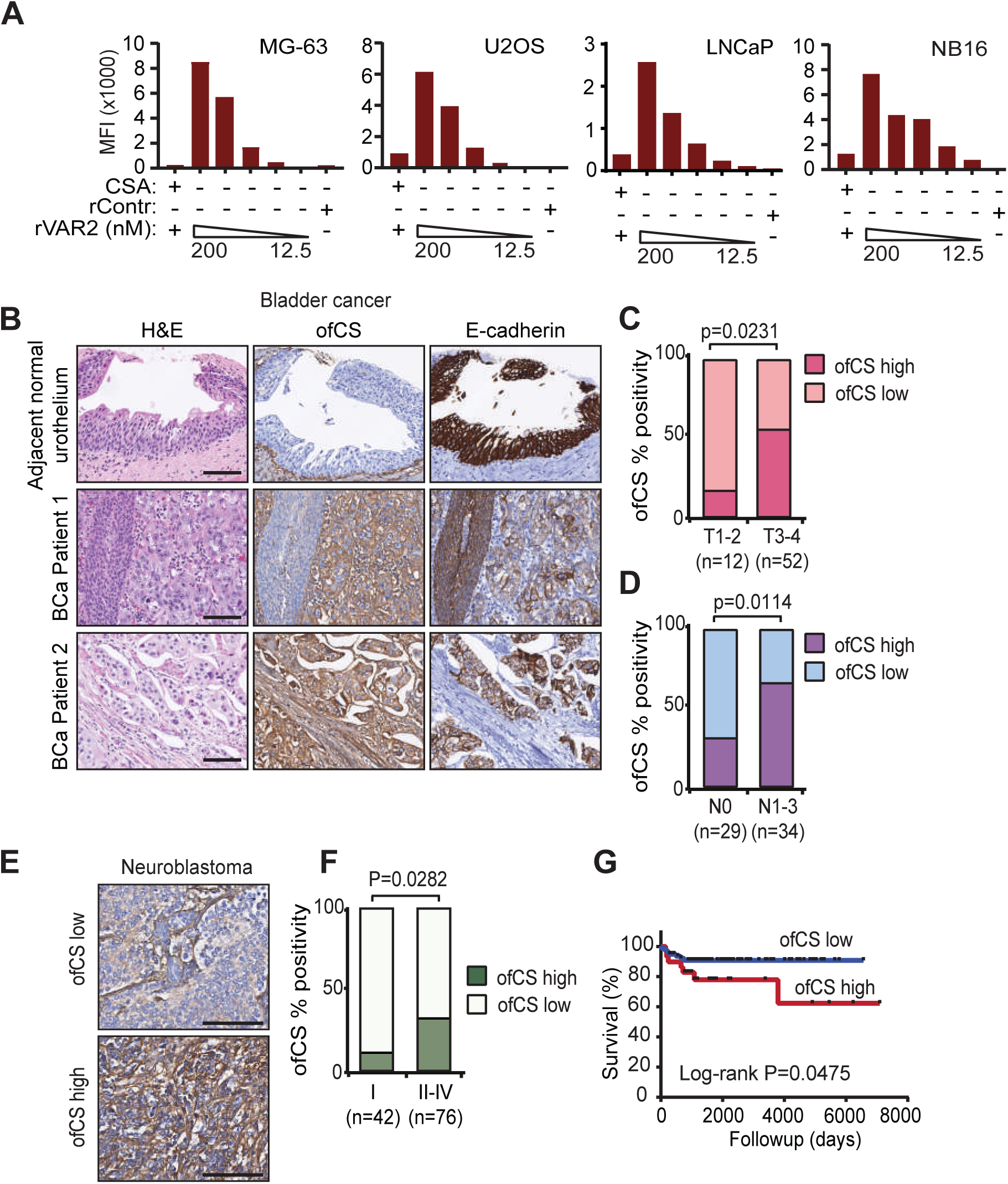
Oncofetal CS expression in solid tumor cell lines and bladder cancer tissue. (**A**) MG-63, U2OS, LNCaP, and NB16 tumor cell lines were incubated with indicated concentrations of rContr protein or VAR2 (12 - 200 nM) +/- purified CSA and analyzed by flow cytometry using anti-V5-FITC. (**B**) Representative H&E and IHC images of normal adjacent urothelium and bladder cancer tissues from two patients. Matched staining images of E-cadherin, as an epithelial marker, in parallel with oncofetal CS (ofCS) detection by VAR2 and anti-V5. (**C**) Bar plot of bladder cancer patient tumors (n=64) indicating ofCS expression in relation to T stage. (**D**) Bar plot of bladder cancer patient tumors (n=63) indicating ofCS expression in relation to N stage. (**E**) Representative IHC images of neuroblastoma tumors selected for high and low oncofetal CS expression. (**F**) Percent ofCS-positive neuroblastoma tumors related to tumor stage. (**G**) Kaplan-Meier plot indicating overall survival of neuroblastoma patients related to oncofetal CS expression. The scale bar represents 100 μm. MIBC: muscle-invasive bladder cancer; oncofetal CS: oncofetal chondroitin sulfate. For statistical analysis in the above panels (C: T stage; D: N stage and F: International Neuroblastoma Staging System (INSS)), two-tailed Fisher’s exact test was used.

**Figure S2.**
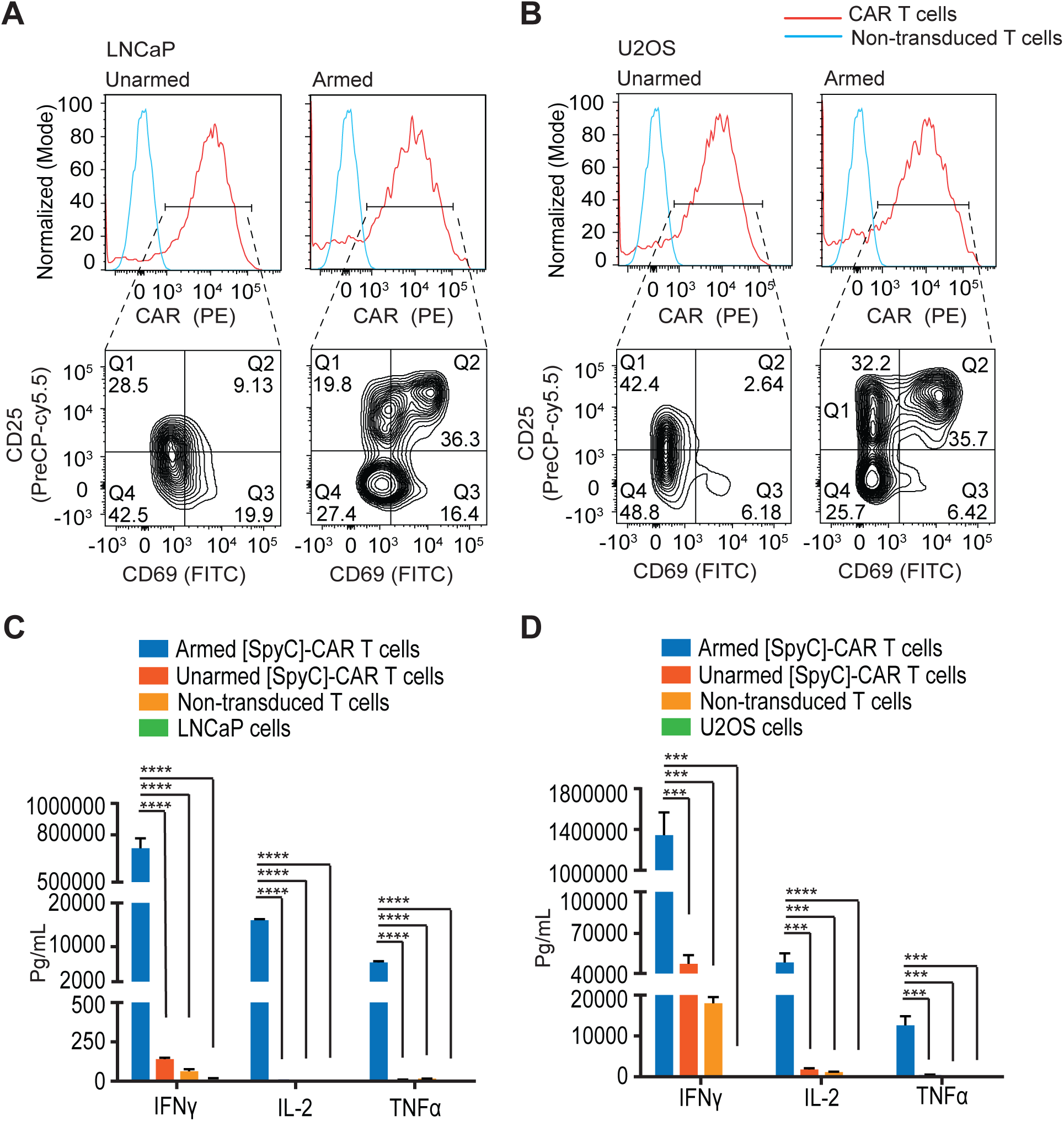
Activation of VAR2-[SpyT][SpyC]-CAR T cells upon tumor cell engagement. (A-B) Armed and unarmed [SpyC]-CAR T cells were incubated with (**A**) LNCaP and (**B**) U2OS cells at a 1:1 E:T ratio for 24 hours before analyzed for Flag, CD69, and CD25 expression by flow cytometry. The results are representative of 3 independent experiments. (C-D) Armed and unarmed [SpyC]-CAR T cells were incubated in triplicate, with (**C**) LNCaP and (**D**) U2OS cells at a 10:1 E:T ratio in 100 μl media, for 48 hours and analyzed for the concentration of indicated cytokines in the culture supernatant. Results are presented as mean ± SEM of three different wells. The statistical significance was determined using one-way ANOVA, Dunnett’s multiple comparison’s test. *, p<0.05; **, p<0.01 ***, p<0.001; ****p<0.0001.

**Figure S3:**
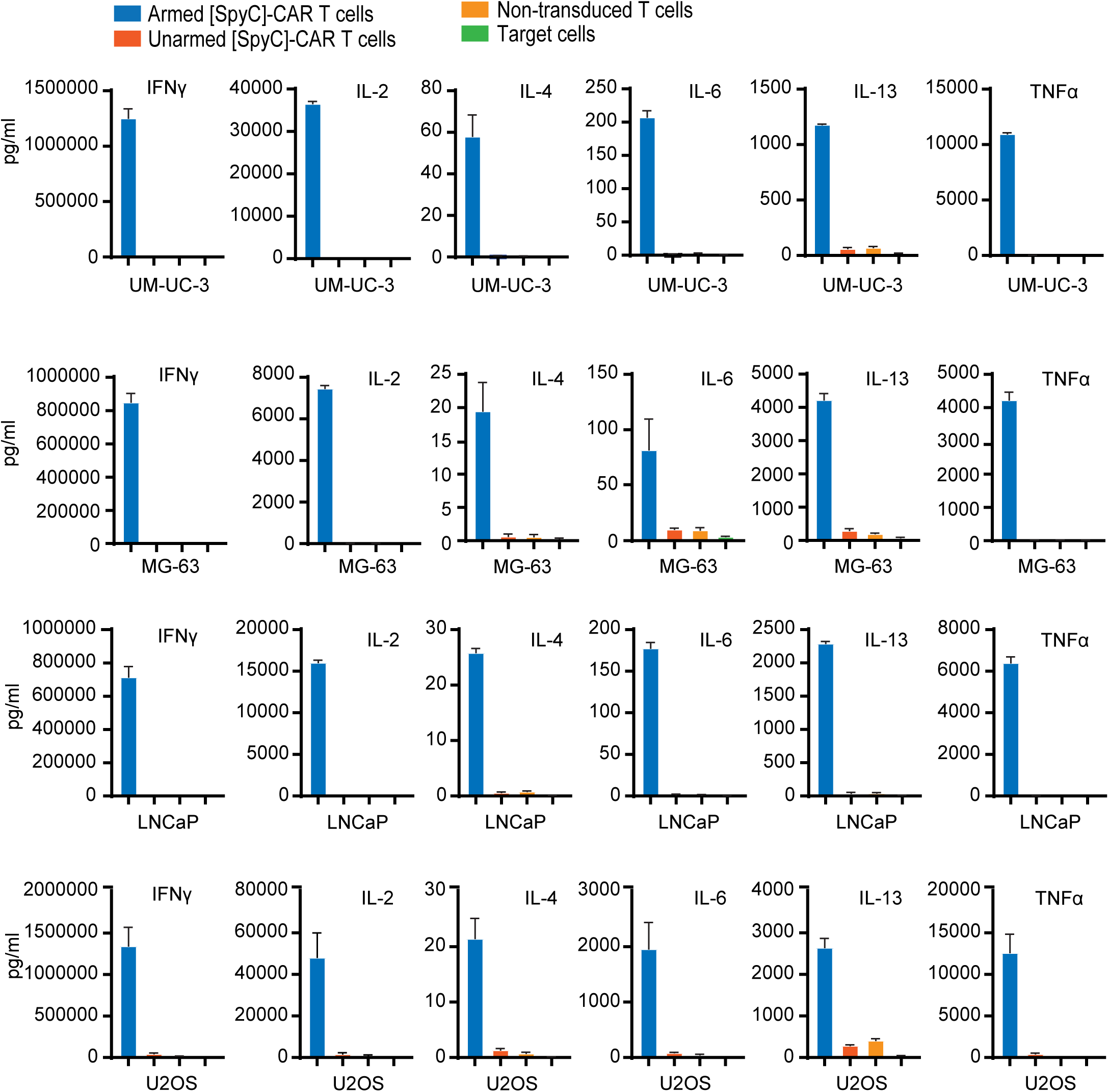
Cytokine responses in co-cultures of effector and target cells. The concentrations of the indicated cytokines in the culture supernatants were assessed after 48 hours of co-culturing between effector T cells formulations and different target cells (i.e., UM-UC3, MG-63, LNCaP, and U2OS), at a 10:1 E:T ratio. Data was analyzed with the Discovery Workbench software. Error bars represent mean ± SEM of three different wells.

**Figure S4:**
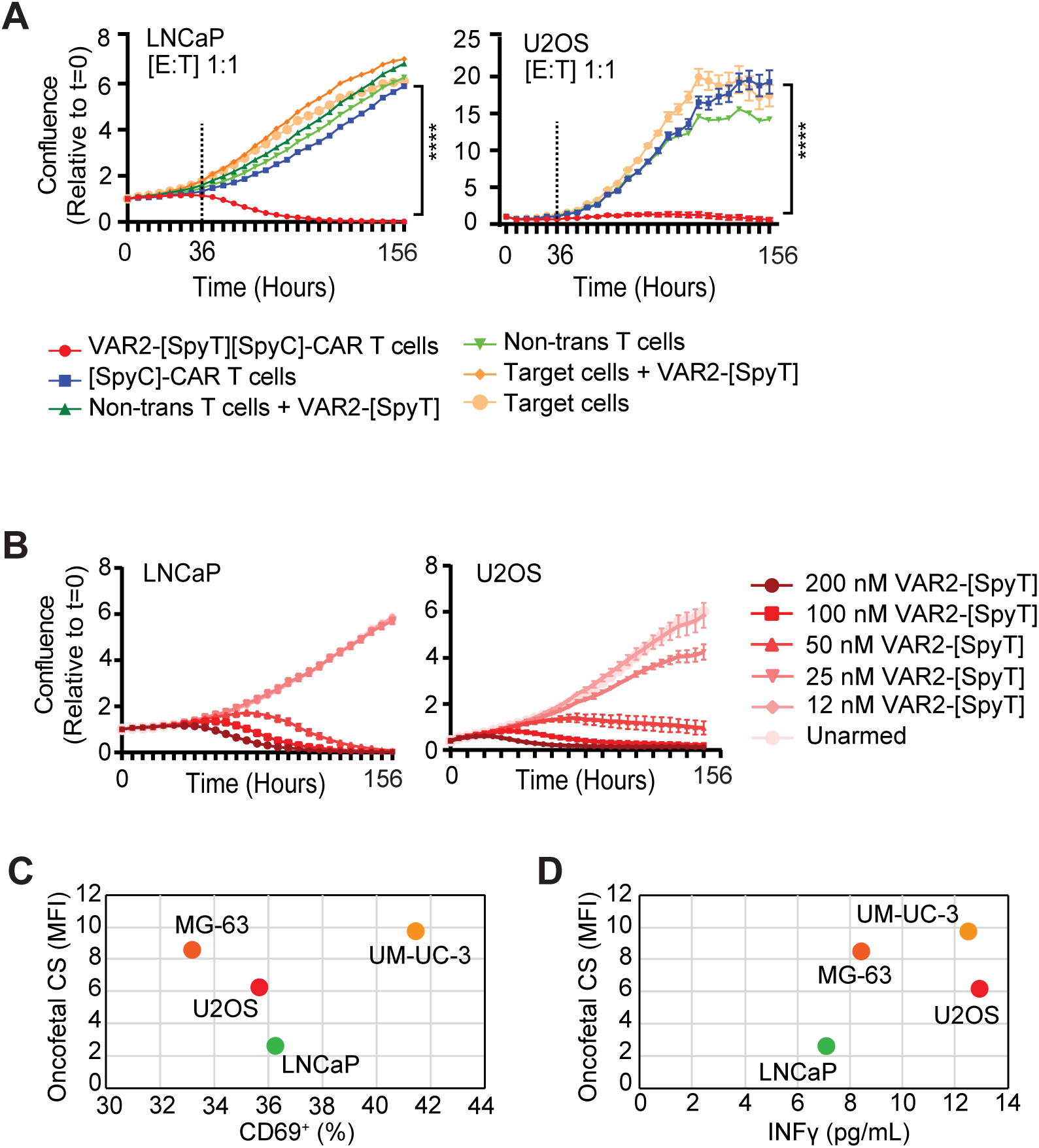
Activity of VAR2-[SpyT][SpyC]-CAR T cells after target cell engagement. (**A**) LNCaP and U2OS target cells (red) were co-cultured in triplicate with indicated formulations of T cells and monitored for one week. Dashed lines indicate time to VAR2-[SpyT][SpyC]-CAR T cell cytotoxicity. Error bars represent mean ± SEM of triplicate wells. Data from a representative donor of three individual donors is shown. Statistics were calculated for the last timepoint, using one-way ANOVA and Dunnett’s multiple comparisons test. (**B**) LNCaP and U2OS target cells (red) were co-cultured with [SpyC]-CAR T cells at a 1:1 E:T ratio with indicated concentrations of VAR2-[SpyT] protein and analyzed for viability using confluence as the readout. Error bars represent mean ± SEM of triplicate wells. (**C**) Percent CD69-positive VAR2-[SpyT][SpyC]-CAR T cells plotted against oncofetal CS expression in indicated target cells. (**D**) IFNγ production (pg/ml) in co-cultures of VAR2-[SpyT][SpyC]-CAR T cells and indicated target cells plotted against oncofetal CS expression of the target cells. All data was analyzed by GraphPad Prism Software.

**Figure S5:**
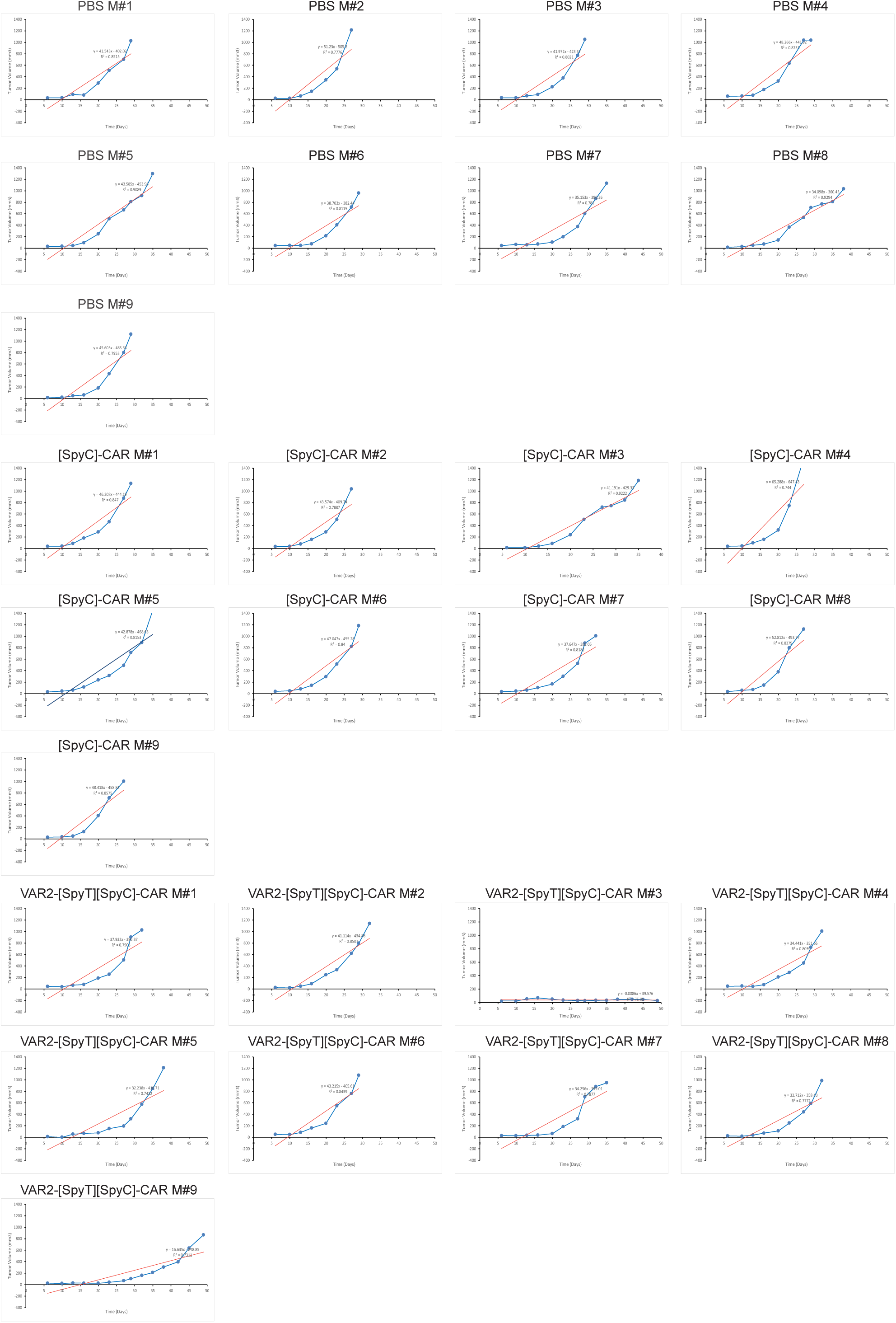
Linear regression analysis of tumor growth. Individual tumor growth curve (blue line) and the slope of the curve (red line) is shown for each mouse treated with PBS, [SpyC]-CAR T cells or VAR2-[SpyT][SpyC]-CAR T cells.

